# Towards Decoding Selective Attention Through Cochlear Implant Electrodes as Sensors in Subjects with Contralateral Acoustic Hearing

**DOI:** 10.1101/2021.08.26.457751

**Authors:** Nina Aldag, Andreas Büchner, Thomas Lenarz, Waldo Nogueira

## Abstract

**Objectives:** Focusing attention on one speaker in a situation with multiple background speakers or noise is referred to as auditory selective attention. Decoding selective attention is an interesting line of research with respect to future brain-guided hearing aids or cochlear implants (CIs) that are designed to adaptively adjust sound processing through cortical feedback loops. This study investigates the feasibility of using the electrodes and backward telemetry of a CI to record electroencephalography (EEG).

**Approach:** The study population included 6 normal-hearing (NH) listeners and 5 CI users with contralateral acoustic hearing. Cortical auditory evoked potentials (CAEP) and selective attention were recorded using a state-of-the-art high-density scalp EEG and, in the case of CI users, also using two CI electrodes as sensors in combination with the backward telemetry system of these devices (iEEG).

**Main results:** The peak amplitudes of the CAEPs recorded with iEEG were lower and the latencies were higher than those recorded with scalp EEG. In the selective attention paradigm with multi-channel scalp EEG the mean decoding accuracy across subjects was 92.0 and 92.5% for NH listeners and CI users, respectively. With single-channel scalp EEG the accuracy decreased to 65.6 and to 75.8% for NH listeners and CI users, respectively, and was above chance level in 9 out of 11 subjects. With the single-channel iEEG, the accuracy for CI users decreased to 70% and was above chance level in 3 out of 5 subjects.

**Significance:** This study shows that single-channel EEG is suitable for auditory selective attention decoding, even though it reduces the decoding quality compared to a multi-channel approach. CI-based iEEG can be used for the purpose of recording CAEPs and decoding selective attention. However, the study also points out the need for further technical development for the CI backward telemetry regarding long-term recordings and the optimal sensor positions.

## 1. Introduction

The cochlear implant (CI) is a neuroprosthesis that is used to rehabilitate severe to profound sensorineural hearing loss. The CI stimulates the auditory nerve electrically and thereby mimics the bottom-up function of the peripheral auditory system. Most CI users obtain good speech recognition in quiet. However, CI users face difficulties to understand speech in the so-called ‘cocktail-party’ scenario (Cherry, 1953), where speech is presented in the presence of simultaneous disturbing sounds (e.g. Bronkhorst, 2000; Zeng et al., 2008).

The ability to focus on a conversation in a noisy ‘cocktail party’ scenario is related to auditory selective attention. The underlying neural mechanisms involved in selective attention are still a topic of current research. The process includes the enhancement and suppression of neural activity related to the ‘auditory object’ (e.g. Alain, Claude and Arnott, 2000) through the ‘bottom-up’ and ‘top-down’ pathway (e.g. Kaya and Elhilali, 2017). Normal-hearing (NH) listeners can easily separate sounds because of their ability to selectively attend to one speaker (e.g. Mesgarani and Chang, 2012). However, CI users cannot fully rely on their selective attention mechanisms, since CIs are designed to mimic the bottom-up auditory pathway and not the top-down neural feedback loop. The idea of decoding selective attention is to reproduce neural selective attention mechanisms through the measurement of neural signals and subsequent signal processing. For this purpose, neural signals are measured through electroencephalography (EEG) or magnetoencephalography (MEG) and filtered in order to reconstruct the attended speech or to decode the locus of attention from the neural signals (e.g. Ding and Simon, 2014; O’Sullivan et al., 2015). The technology may find application in closed-loop CI or hearing aid systems, e.g. for neurally-steered hearing prostheses that can identify the attended speech to steer speech enhancement algorithms (e.g. O’Sullivan et al., 2017). Other applications can be found in the objective fitting of CIs or in the clinical evaluation of hearing development in children or disabled CI users (e.g. Finke et al., 2017; Mc Laughlin et al., 2013).

The feasibility of decoding selective attention from single-trial EEG has been shown in various previous studies in NH listeners (e.g. Mirkovic et al., 2015; O’Sullivan et al., 2015) and CI users (Nogueira et al., 2019a, 2019b; Paul et al., 2020). The use of scalp EEG, however, is not applicable in the clinical routine or in everyday applications. Around-the-ear-EEG, as used by Bleichner et al. (2016) and Mirkovic et al. (2016) in NH listeners and by Nogueira et al. (2019b) in CI users, and in-ear-EEG, as used by Fiedler et al. (2017) in NH listeners, is one possibility to reduce the visibility and to maximize the portability of the recording system. However, non-invasive recordings from the scalp can contain high artifact, caused by electrode displacement, body motion, ocular and muscle movement (Islam et al., 2016). The quality of invasive EEG is often better because invasive EEG is less prone to ocular artifacts (Ball et al., 2009), filter effects of the tissue and spatial distortions from the skull (Waldert, 2016).

Modern CIs are equipped with backward telemetry capabilities that can be used to record voltage fluctuations around the electrode contacts. The idea of this work was, to use electrodes of a CI in combination with the CI backward telemetry system to record neural signals and to decode selective attention from these CI-based recordings. The CI backward telemetry systems are primarily used for impedance measurement, electrocochleography (ECochG) and the recording of electrically evoked compound action potentials (ECAPs). ECochG can be used to actively monitor cochlear responses during the electrode insertion to assist in predicting electrode location and preventing intraoperative trauma (e.g. Acharya et al., 2016; Calloway et al., 2014; Harris et al., 2017), to estimate residual hearing objectively (e.g. Koka et al., 2017) or to characterize electric and acoustic interaction (e.g. Imsiecke et al., 2020; Krüger et al., 2020). The recording of ECAPs is an established method to assess the CI function and the auditory nerve function intra- and postoperatively (e.g. Christov et al., 2016) and may assist in the fitting of CIs (e.g. Botros and Psarros, 2010; McKay et al., 2013; McKay and Smale, 2017). ECAPs and ECochG have been successfully transferred into clinical applications. Similarly, the advantages of recording EEG with invasive CI electrodes would be the easy applicability in CI users and the proximity between the electrodes and the auditory cortex. However, most of the currently available CI backward telemetry is limited by the short measurement windows due to hardware restrictions. McLaughlin et al. (2012) used the telemetry system of CIs from the company Cochlear Ltd. to measure early AEPs, also known as the auditory brainstem responses (ABR), and late AEPs, also known as cortical auditory evoked potentials (CAEPs). Recording of CAEPs, which normally requires continuous recording of up to 1 s, was realized by a combination of repeated measurements with shifting time windows. This procedure was highly time consuming and thus is not practically applicable in clinical routine measurements. Somers et al. (2021) performed continuous EEG recording via a CI from the company Cochlear Ltd. using a percutaneous connector. Moreover, a high-quality external amplifier could be used and the invasive CI electrodes could be combined with non-invasive scalp channels for a hybrid EEG recording. However, Somers et al. (2021) approach also lacks the practical applicability because it requires an invasive procedure which goes in hand with an increase of multiple risks for the patient, such as infection. Nevertheless, both approaches prove the general feasibility of CI-based recording of cortical signals such as CAEPs and motivate the development of fully implantable CI-based EEG (iEEG).

In the current study, the iEEG was based on the electrodes and the backward telemetry system of Advanced Bionics CIs. With the current system it is not possible to record and to stimulate simultaneously. For this reason, the study population included CI users with contralateral acoustic hearing, so that it was possible to record cortical signals on the CI side while presenting the stimuli on the acoustically hearing ear. This study population will be referred to as CI users along this manuscript. First, CAEPs were measured to investigate the general feasibility of recording cortical potentials with the CI. Another restriction of the current system was that only a single recording channel was utilizable. For this reason, it was investigated if selective attention decoding was possible from single-channel recordings and with monaural stimulus presentation. Those single-channel scalp EEG recordings were compared to the single-channel iEEG using a selective attention paradigm. The results of these experiments offer insights into the current restrictions and possibilities and might promote further optimization of CI-based recording systems.

## 2. Methods

### 2.1 Subjects

5 CI users with contralateral acoustic hearing (4 male, 1 female; mean age: 66 years) and 6 normal hearing (NH) listeners (5 male, 1 female; mean age: 33 years) participated in the study. All subjects were native German speakers and reported no history of neurological disorders. The demographics of NH listeners and CI users and their measurement conditions are provided in Table 1 and 2, respectively.

**Table 1:**
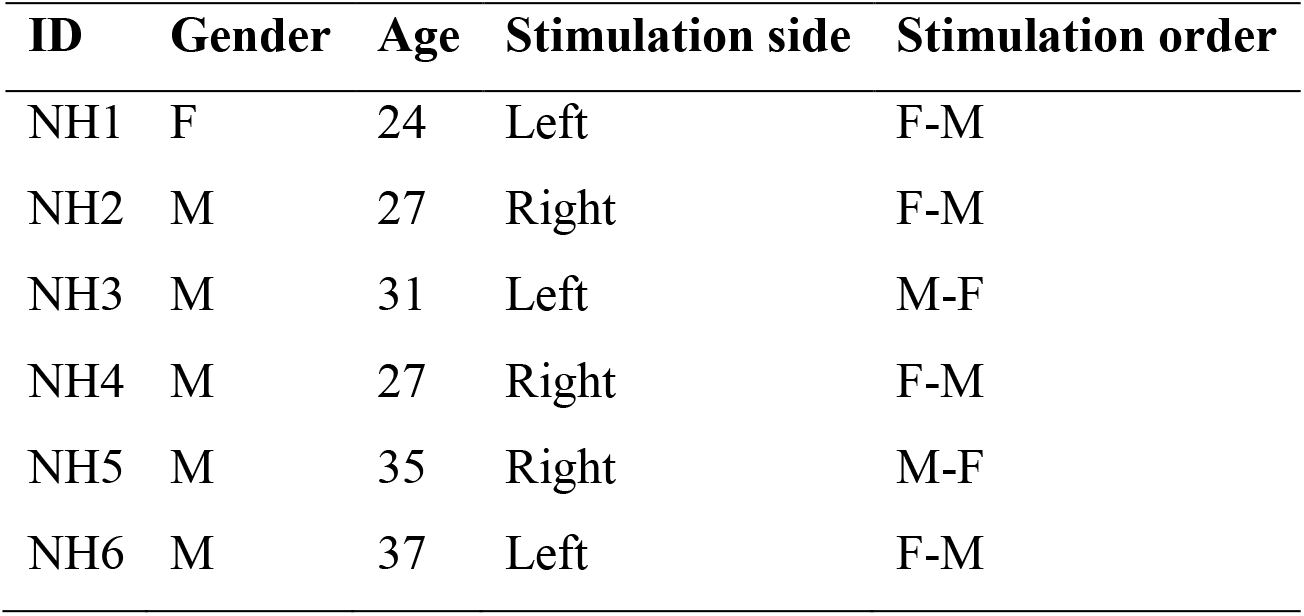
Demographics of normal hearing (NH) listeners. F = female; M = male.

**Table 2:** Demographics of cochlear implant (CI) users with contralateral acoustic hearing. F = female; M = male.

| ID | Gender | Age | Implant | Electrode | Stimulation<br>side | Stimulation<br>order |
| --- | --- | --- | --- | --- | --- | --- |
| CI1 | M | 73 | Ultra 3D | Mid-Scala | Left | M-F |
| CI2 | M | 73 | HiRes90K Advantage | Mid-Scala | Right | F-M |
| CI3 | M | 53 | Ultra | Mid-Scala | Left | M-F |
| CI4 | M | 80 | Ultra | Mid-Scala | Left | F-M |
| CI5 | F | 49 | Ultra 3D | SlimJ | Left | M-F |

The CI users were unilaterally implanted with an Advanced Bionics device with a fully inserted electrode array. All CI users had residual acoustic hearing on the contralateral ear to the CI. The audiograms, measured on the acoustically hearing ear only, are provided in Figure 1. Subject CI1, CI3 and CI4 wore a hearing aid on the acoustically hearing ear in daily life. The NH listeners served as a control group. All NH listeners had age-appropriate hearing with a hearing loss equal to or lower than 10 dB hearing level in the frequency range from 0.25 to 8 kHz.

**Figure 1:**
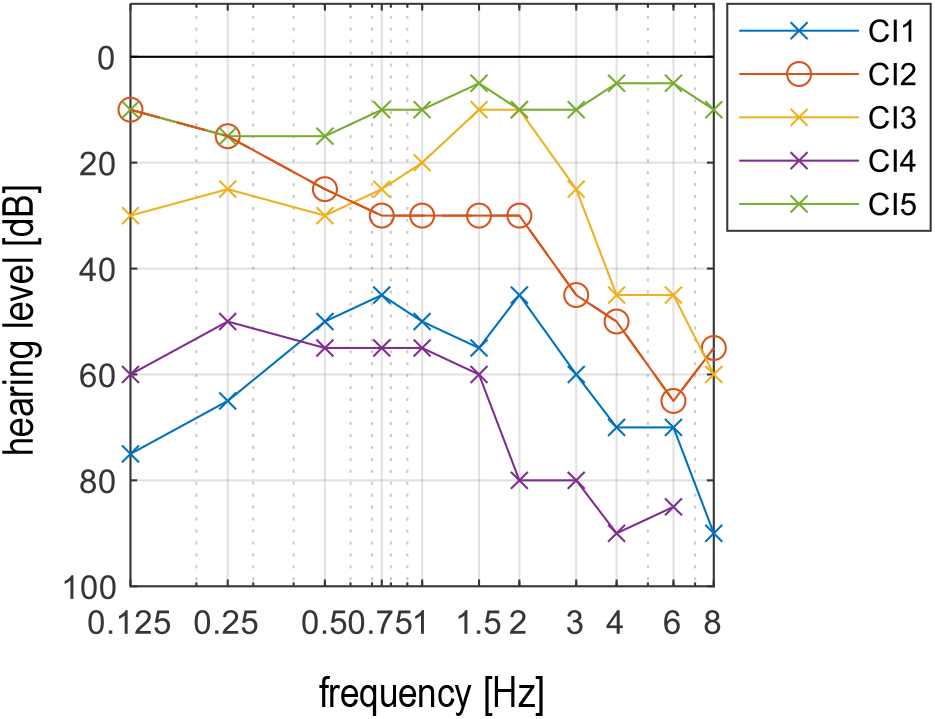
Audiograms of cochlear implant (CI) users on the acoustically hearing ear contralateral to the CI.

### 2.2 Experimental Setup

The experimental setup is shown in Figure 2. The current study extended the experimental setup from Nogueira et al. (2019a and 2019b) to record iEEG through the backward telemetry and the electrodes of a CI.

**Figure 2:**
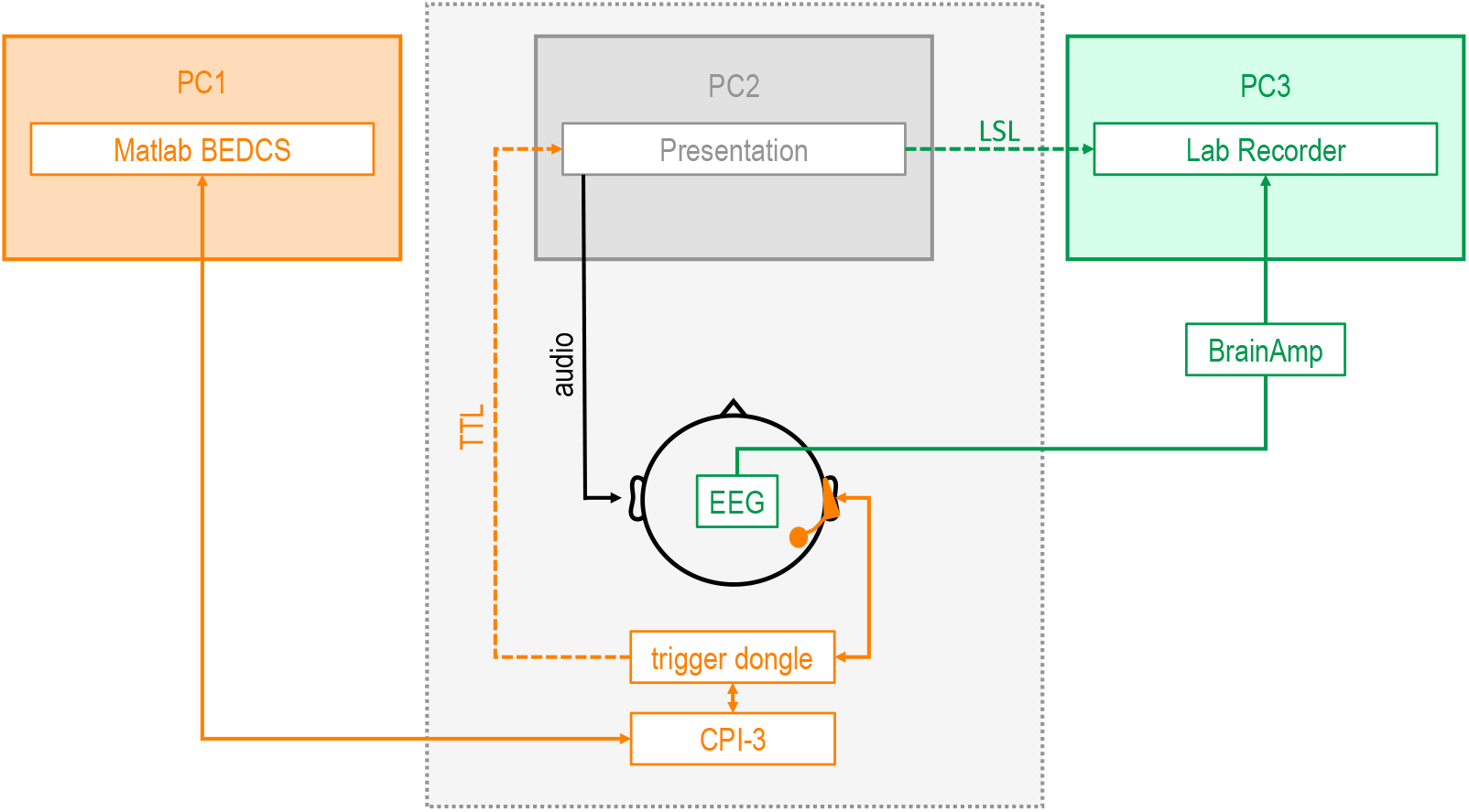
Experimental Setup: The subject was sitting in an electromagnetically and acoustically shielded chamber. Acoustic stimuli were presented via in-ear phones in the contralateral ear to the cochlear implant (CI) and optical input was provided via a screen. The scalp electroencephalography (EEG) recording (green) was synchronized through event markers in lab streaming layer (LSL) between personal computer 2 and 3 (PC2, PC3). The recording via CI (orange) was captured by PC1 and synchronized through transistor-transistor logic (TTL) triggering between the trigger dongle and PC2. BEDCS = bionic ear data collection system; CPI = clinical programming interface.

Subjects were sitting in an electromagnetically and acoustically shielded chamber. The software Presentation version 20.1 (Neurobehavioral Systems, Inc., Berkeley, California) was used to present acoustic stimuli and visual input via a screen. Acoustic stimuli were presented via an amplification system (A-10, Pioneer Corporation, Tokyo, Japan) and inserted earphones (E-A-RTONE Gold 3A, 3M, St. Paul, Minneapolis).

The scalp EEG system consisted of an infracerebral electrode cap (Easycap GmbH, Herrschingen, Germany) and a BrainAmp amplification system (Brain Products GmbH, Gilchingen, Germany). The electrode cap comprised 96 passive sintered silver/silver chloride electrodes. A reference electrode was placed on the nose tip and a ground electrode on the forehead anterior to the midline frontal electrode (Fz). Recordings were performed with a sampling frequency of 1 kHz and an online filter from 0.2 to 250 Hz. The data was stored by Lab Recorder (Neurobehavioral Systems, Inc., Berkeley, California) in an extensible data format (.xdf).

For the iEEG the subject’s individual internal unit of the CI was connected to an external unit through a coil and a Naida Q90 sound processor. A custom made trigger dongle and a clinical programming interface (CPI 3; Advanced Bionics AG, Stäfa, Switzerland) connected the CI via USB to the research interface bionic ear data collection system (BEDCS) app version 3.1 (Advanced Bionics AG, Stäfa, Switzerland), implemented in Matlab 2019b (The MathWorks, Inc., Natick, Massachusetts). The data was stored in a Matlab file format (.mat). The equipment for the iEEG was provided by Advanced Bionics LLC (Valencia, United States). The iEEG recordings were performed with a sampling frequency of 5625 Hz. Single-channel recordings were performed between the most apical intracochlear electrode (E0) as the active input and an extracochlear electrode, located on the implant case, as the reference input. An overview on the electrode positions is provided in Figure 3.

**Figure 3:**
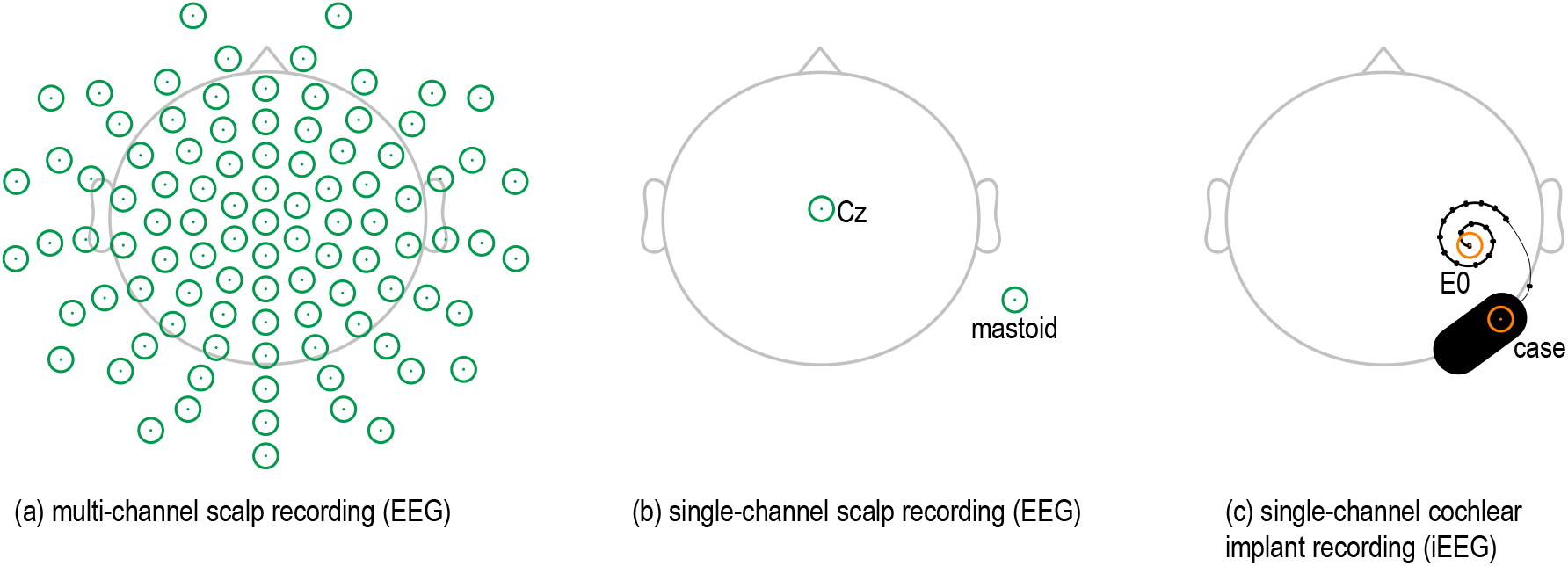
Electrode configurations. (a) All scalp electroencephalography (EEG) electrodes were used, resulting in a 96-channel recording. (b) The vertex electrode (Cz) was re-referenced against the electrode on the mastoid contralateral to the ear that was presented with the acoustic stimuli, resulting in a single-channel recording. (c) Two cochlear implant electrodes were used, resulting in a single-channel implant-based EEG (iEEG). The most apical intracochlear electrode (E0) was the active input and the extracochlear implant case electrode was the reference input.

### 2.3 Stimuli

CAEPs were evoked through linearly-ramped 1 kHz sine tone burst stimuli. The rise and fall time of the ramp was 10 ms and the plateau time was 30 ms. The stimuli were presented 100 times with a repetition rate of ∼1 Hz. The measurement was conducted with three different loudness levels. The subject’s individual loudness was adjusted through a ten-point loudness-rating scale (with 1 equivalent to ‘extremely soft’ and 10 equivalent to ‘extremely loud’). The first level was defined as 7 (‘loud but comfortable’) and the second level as 3 (‘soft’). The third level was without stimulus presentation and served as a baseline recording.

The current study used the same selective attention paradigm as in Nogueira et al. (2019a, 2019b). Two German stories, ‘A drama in the air’ by Jules Verne and ‘The two brothers’ by the Grimm Brothers, were presented to the subject. ‘A drama in the air’ was narrated by a male speaker, ‘The two brothers’ was narrated by a female speaker. In contrast to the two previous studies, the two stories were digitally combined and presented through one acoustic ear. For all bimodal listeners, the stories were pre-processed digitally to amplify the sounds based on their individual audiogram. This was done by applying the ‘Half-Gain Rule’, as e.g. in Krüger et al. (2020). The loudness was adjusted by the subject to reach level 6 (‘comfortable’) using the ten-point loudness-rating described above.

The stories were 24 min long and were divided into three sections of 8 min. Every section was divided into four subblocks of 2 min. After each section, the subject was able to take a break. The stories were presented twice, with the subject attending to one story the first time and to the other story the second time. At the start of each subblock, the subject was instructed to attend to a particular voice. The order of the to-be-attended stories was randomized across subjects (see Tables 1 and 2 for details). Altogether, the procedure included 48 min recording time for each subject, excluding the time for breaks.

### 2.4 Behavioral Response

In the selective attention experiment the subject was asked to answer questions about the attended story after each subblock to make sure that the correct story was attended. 4 single-choice questions with 4 possible answers were displayed on the screen. The questions were answered with the numbers 1 to 4 on a keyboard.

At the end of each story the subject was asked 3 questions about the difficulty of the task: “How difficult was the task in general?”, “How easily could you focus on the story?” and “How much did the other story distract you?”. The answers were rated from 1 to 4, with 1 equivalent to ‘very easy’ or ‘not at all’ and 4 equivalent to ‘very difficult’ or ‘very much’.

### 2.5 Test Procedure

A pure-tone audiogram was measured to determine the hearing status of the acoustic ear (CI users) and to validate NH status in the reference group. The subject’s hair was washed and dried and the EEG cap was mounted onto the subjects head. The impedances of the scalp EEG electrodes were maintained below 20 kΩ by cleaning the skin with ethanol and applying an abrasive and conductive electrolyte gel (Abralyt 2000; Easycap GmbH, Herrschingen, Germany). CAEPs and selective attention measurements were recorded. During both measurements, the subject was asked to sit calm and relaxed in the measurement chamber, to keep the eyes open and to keep the visual focus on a white fixation cross on the black screen in front of the subject. Regular breaks were provided according to the needs of the subject. All sources of electromagnetic interference were removed from the chamber and the light was dimmed.

### 2.5 Data Analysis

#### 2.5.1 Pre-Processing

Pre-processing of EEG data was performed offline in Matlab 2020a (The MathWorks, Inc., Natick, Massachusetts) using the EEGLAB toolbox (v2020.0; Delorme and Makeig, 2004). The constant time delays of the EEG systems were compensated, i.e. the EEG had a delay of 77 ms with respect to the acoustic stimulation (Nogueira et al., 2019b) and the iEEG started 250 ms before the acoustic stimulus was presented. A drift in iEEG data was removed by subtracting the low-pass filtered signal from the original recording. The low-pass filter was based on a moving average filter with a window width of 200 ms.

CAEP iEEG and EEG data was filtered with a 4^th^ order Butterworth bandpass filter from 1 to 20 Hz, using the filtfilt-function in Matlab for zero-phase filtering, and downsampled to 500 Hz. The data were epoched in 100 trials. The first 5 trials were excluded and the mean of all other trials was calculated. Significant peaks were identified using bootstrapping with 200 replicates.

Selective attention iEEG and EEG data was pre-processed according to the procedure of Nogueira et al. (2019a and 2019b). EEG and iEEG were filtered using a 4^th^ order Butterworth bandpass filter from 2 to 8 Hz, using the filtfilt-function in Matlab for zero-phase filtering, and downsampled to 64 Hz. The EEG data was epoched in 48 trials of 60 s for each subject. The drift in iEEG data caused a signal saturation after a variable amount of time between 15 and 90 s, depending on the individual CI and the radio-frequency link. The parts of the signal that were saturated were discarded, leading to a reduced amount of valid iEEG data. The amount of valid iEEG data was 36 min for subjects CI1 and CI2, 6 min for CI3 and CI4 and 18 min for CI5. The iEEG data was concatenated to trials of 60 s.

The envelope **x**[k] of the audio signal was calculated as the absolute value of the complex analytic signal that is composed of the signal ***s*** as the real part and its Hilbert transformation ***s***_H_ as the complex part:

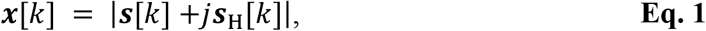

with *k* = 0 … *K* − 1 being the time samples. The audio signal was low-pass filtered below 8 Hz using a 4^th^ order Butterworth low-pass filter with a cutoff frequency of 8 Hz with the Matlab filtfilt-function and downsampled to 64 Hz.

#### 2.5.2 Selective Attention Decoding Architecture

Linear forward models assume that the EEG represents a convolution of the speech envelope with an impulse response, referred to as temporal response function (TRF). The multivariate TRF (mTRF) toolbox, provided by Crosse et al. (2016), was used to investigate how the speech envelope is encoded in the brain.

The TRF is represented as the encoder weights of the forward model ***W***_f,*n*_. ***W***_f,*n*_ was estimated through least squares error minimization between the encoded neural response ***ŷ***_*n*_[*k*] and the recorded neural response ***y***_*n*_[*k*] at each time sample *k* = 0 … *K* − 1 and for each channel *n* = 0 … *N* − 1. Tikhonov regularization was applied to avoid overfitting, with *λ* as the regularization parameter:

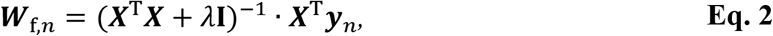

whereby **I** is the identity matrix and ***X*** is a *K* × *L* matrix of lagged speech envelopes. The lag window *l* = 0 … *L* − 1 was used to include temporal context. ***X***_a_ contains the attended and ***X***_u_ contains the unattended lagged speech envelopes, respectively:

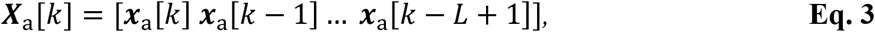

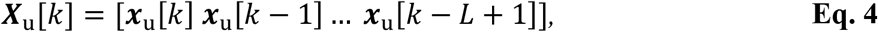

The disadvantage of forward models is that these represent an independent univariate filter for each channel *n*. In contrast, backward models treat the channels in a multivariate context and are therefore superior regarding the reconstruction performance (Crosse et al., 2016). The reconstruction of the attended speech stream was accomplished through backward modelling using the mTRF toolbox. We followed the same procedure as previous works (e.g. Mirkovic et al., 2015; Nogueira et al., 2019a, 2019b; O’Sullivan et al., 2015) in which the envelope of the attended audio signal was reconstructed with a linear spatio-temporal filter, also named decoder. The envelope of the attended speech signal ***x***_a_[*k*] was reconstructed as follows:

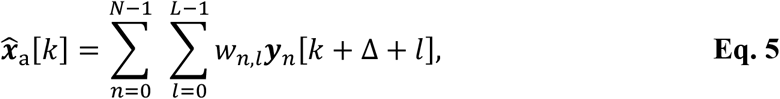

whereby 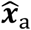 is the envelope reconstruction, *w*_*n,l*_ is the decoder weight for each electrode and sample and ***y***_*n*_ is the recorded neural response at each electrode. A lag Δ between the envelope and the neural response was introduced to model the response latency of the auditory pathway.

The decoder weights of the backward model, *w*_*n,l*_ or ***W***_b_ in vector notation, were estimated through least squares error minimization between the reconstruction 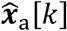 and the original attended speech envelope ***x***_a_[*k*] using Tikhonov regularization:

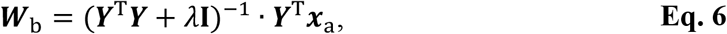

whereby **I** is the identity matrix and ***Y*** is a *K* × *NL* matrix of lagged neural responses. For a single-channel signal ***Y***_*n*_ is defined in Eq. 7, while for a multi-channel signal each column of ***Y*** is replaced by N columns (see Crosse et al. (2016) for details):

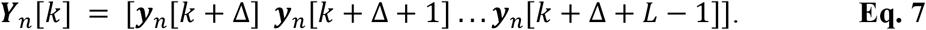

The reconstructed signals 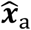 were classified into correct and incorrect trials based on the Pearson correlation coefficient *ρ*. If *ρ*_a_ between 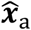 and ***x***_a_ was higher than *ρ*_u_ between 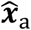 and the envelope of the unattended speech signal ***x***_u_ a trial was defined as correct. The decoding accuracy was defined as the percentage of correctly classified trials. The full reconstruction and decision process is shown in Figure 4.

**Figure 4:**
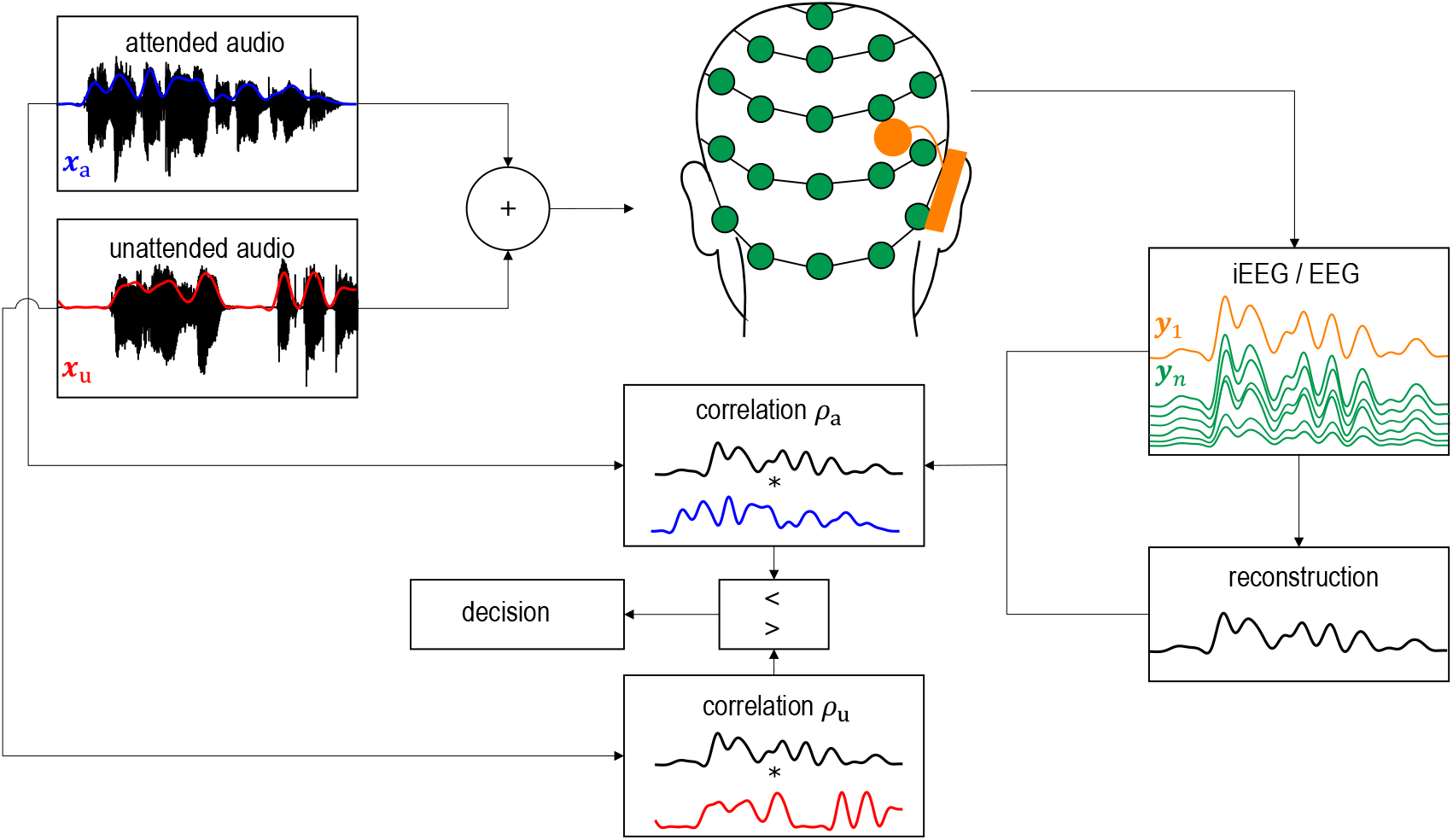
Reconstruction-based architecture that was used to decode selective attention. The envelope of the attended speech stream was reconstructed from the electroencephalography (EEG, green) or implant-based EEG (iEEG, orange) by means of linear backward decoding. The model classified correct and incorrect decisions based on the Pearson correlation coefficient between the reconstructed envelope and the envelopes of the attended (blue) and unattended (red) speech stream, respectively.

***W***_f_ and ***W***_b_ were determined in a training process with 60 s trials (3840 samples). The regularization parameter *λ* that resulted in highest decoding accuracies was empirically determined. The TRF ***W***_f_ was evaluated for a lag window of *L* = 33, including temporal context from 0 to 500 ms pre-stimulus, to investigate the neural ressponse to the stimuli. The decoder ***W***_b_ was evaluated repeatedly for 33 non-overlapping lag windows of 15.6 ms (*L* = 2), covering a total period from 0 to 480 ms post-stimulus, to investigate the optimal lag Δ of the recorded EEG with respect to the speech envelope.

The decoding accuracy was evaluated using leave-one-out cross-validation. The chance level was determined using a binomial test at 5 % significance level (Combrisson and Jerbi, 2015). For EEG the chance level was 38.6 – 61.4 %, corresponding to 24 trials per class. For iEEG the chance level depended on the amount of data that was available for each subject: CI1 and CI2 (18 trials per class: 37.0 – 63.0 %,), CI3 and CI4 (3 trials per class: 24.0 – 76.0 %) and CI5 (9 trials per class: 32.5 – 67.5 %).

## 3. Results

### 3.1 Cortical Auditory Evoked Potentials (CAEPs)

The CAEPs for all CI users are shown in Figure 5. Peaks that were significantly above baseline level were identified using bootstrapping with 200 replications. Significant peaks were labeled as N1 (80 to 150 ms) and P2 (150 to 250 ms), according to their latency with respect to the stimulus. The characteristic morphology of the peaks was most clear in the scalp EEG, using the vertex electrode Cz, re-referenced to the mastoid contralateral to the stimulus. Significant N1 and P2 in EEG were detected for subjects CI2, CI4 and CI5. For subject CI1 only a significant N1 was detected and for subject CI3 no significant peaks were detected at all. Peaks were consistent across the two stimulus levels, with the exception of subject CI5.

**Figure 5:**
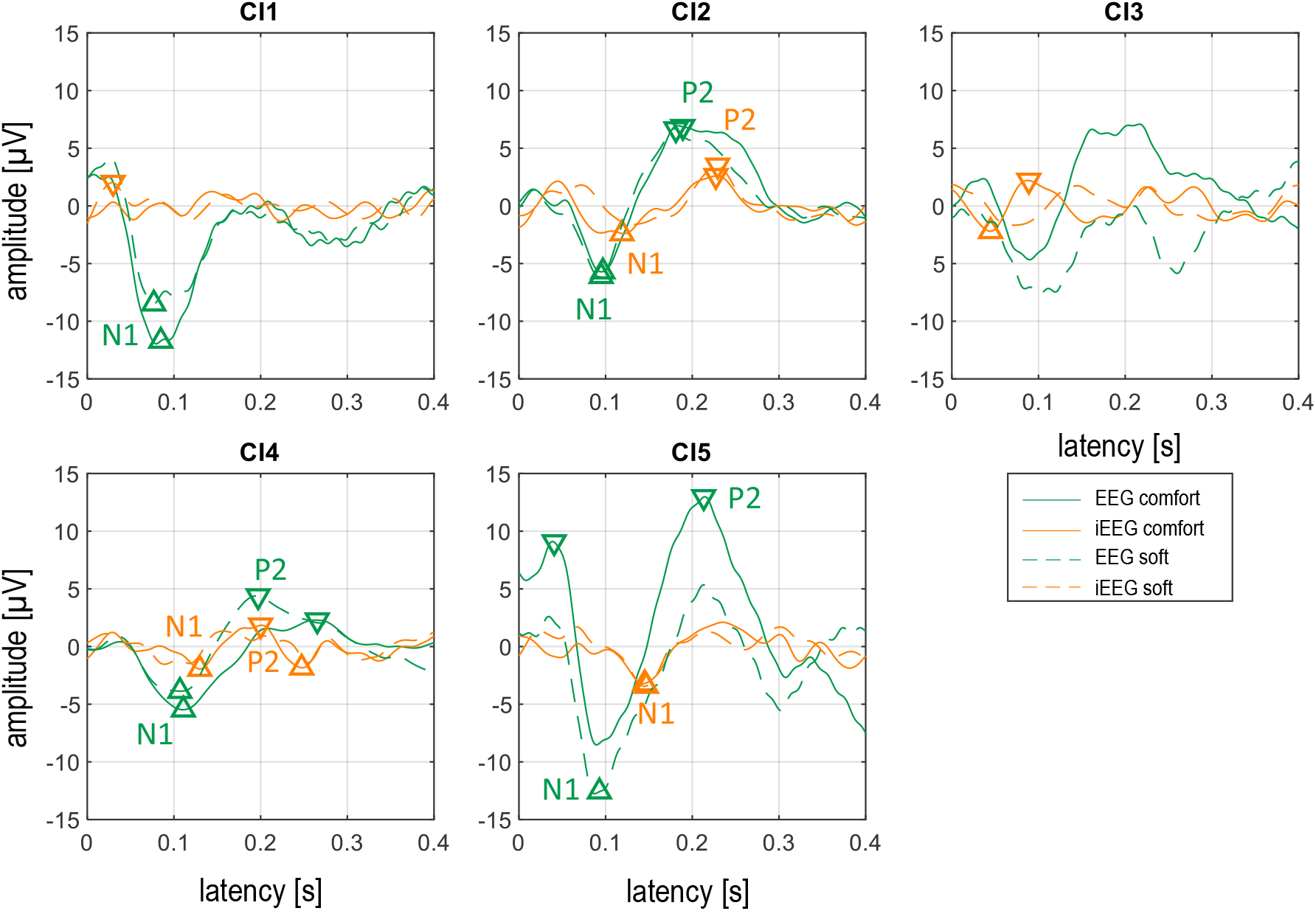
Cortical auditory evoked potentials (CAEPs) for all cochlear implant users (CI1-CI5) measured with scalp electroencephalography (EEG; Cz - contralateral mastoid; green lines) and with implant-based EEG (iEEG; case - E0; orange lines) as a response to ‘loud but comfortable’ (solid lines) and to ‘soft’ stimuli (dashed lines). Significant peaks were marked and additionally labeled when they were in the range of typical N1 or P2 peaks.

The N1 and P2 responses recorded with iEEG had smaller amplitudes and higher latencies than the responses recorded with EEG. Significant N1 and P2 peaks in iEEG were detected for subjects CI2 and CI4. For subject CI5 only a significant N1 was detected and for subjects CI1 and CI3 no significant N1 or P2 peaks were detected at all. Peaks were consistent across the two stimulus levels for subject CI2 and CI5, while for subject CI4 the significant peaks could only be observed for the ‘loud but comfortable’ stimulation level.

### 3.2 Selective Attention Behavioral Data

The percentage of correctly answered questions during the breaks of the selective attention experiment was calculated for every subject. A chance level of 16.8 to 33.2 % was determined using a binomial test at 5 % significance level (Combrisson and Jerbi, 2015). All subjects performed above chance level (mean = 77 %, σ = 19 %). On average, NH listeners and CI users obtained 87 % and 66 % correct responses, respectively.

The difficulty of the task was rated with an average score of 2.2 (σ = 0.5), which was equivalent to ‘rather easy’. The story narrated by a female voice was reported to be easier (mean score = 1.9) than the story narrated by a male voice (mean score = 2.5). On average, NH listeners rated the difficulty of the task to be lower (mean score = 2.0) than CI users (mean score = 2.5).

### 3.3 Selective Attention Temporal Response Function (TRF)

The TRF is an indicator of how well the acoustic speech envelope is encoded in the neural data (Haufe et al., 2014). The TRF was calculated using the linear forward model for NH listeners and CI users based on scalp EEG (Figure 6). The morphology of the TRF weights across lags is very similar to the typical morphology of CAEPs across latencies, showing a negative peak at around 100 ms and a positive peak at around 200 ms. The TRF that encodes the attended speech envelope presents higher weights than the TRF that encodes the unattended speech envelope. Moreover, the TRF weights present highest values in areas around the vertex.

**Figure 6:**
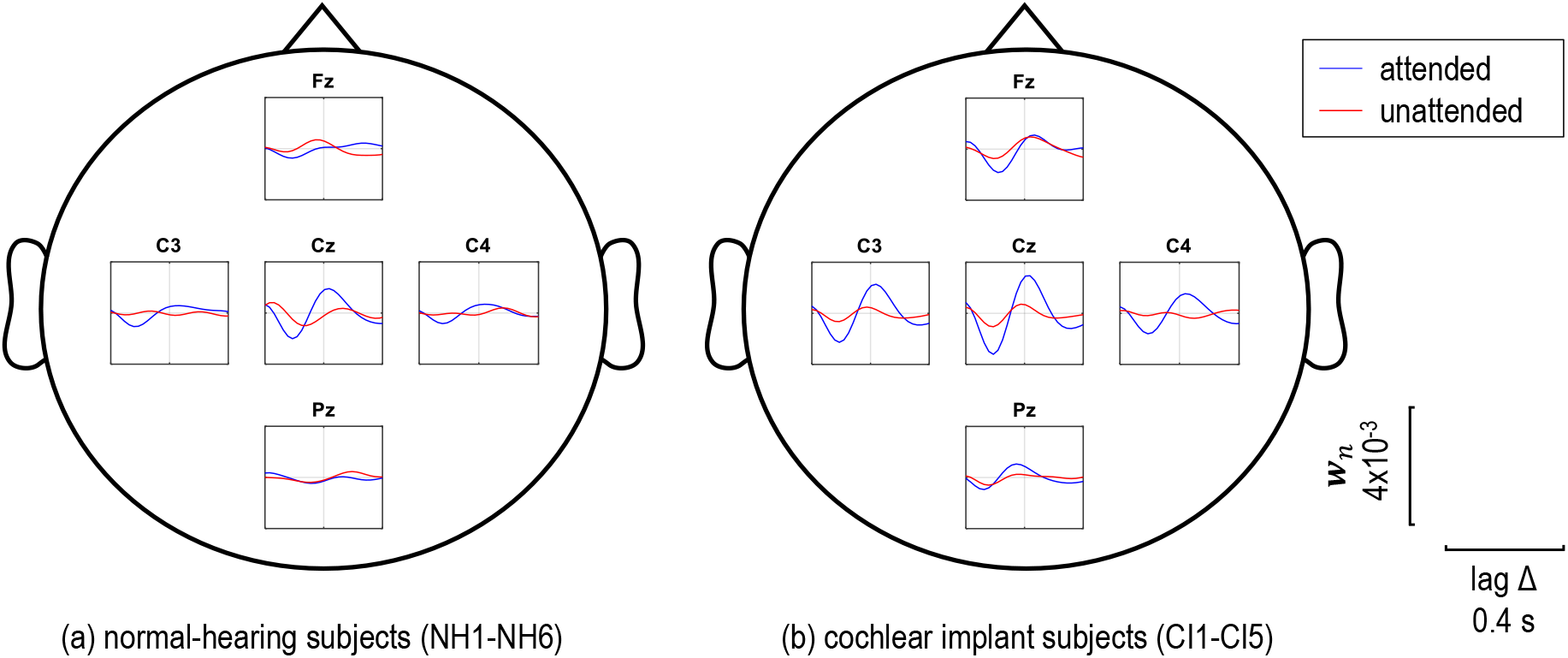
Mean temporal response functions (TRFs) recorded through scalp electroencephalography (EEG) of (a) normal-hearing listeners (NH1-NH6) and (b) cochlear implant users (CI1-CI5). The linear forward model was trained with the envelope of the attended speech (blue) or the envelope of the unattended speech (red). The encoder weights ***w***_*n*_ for 5 representative electrodes (Fz, C3, Cz, C4 and Pz), re-referenced against the common average, are shown across lag Δ between stimulus and recording.

The TFR based on iEEG was calculated for each CI user individually (Figure 7). As for CAEPs, the TRF shows a high inter-subject variability. For subjects CI1 and CI3, the peaks of the attended TRF correspond to the N1-P2 morphology of a typical CAEP. For subjects CI2, CI4 and CI5, the peaks are inverted in amplitude. In addition, later peaks after 200 ms occur in the TRF of all subjects.

**Figure 7:**
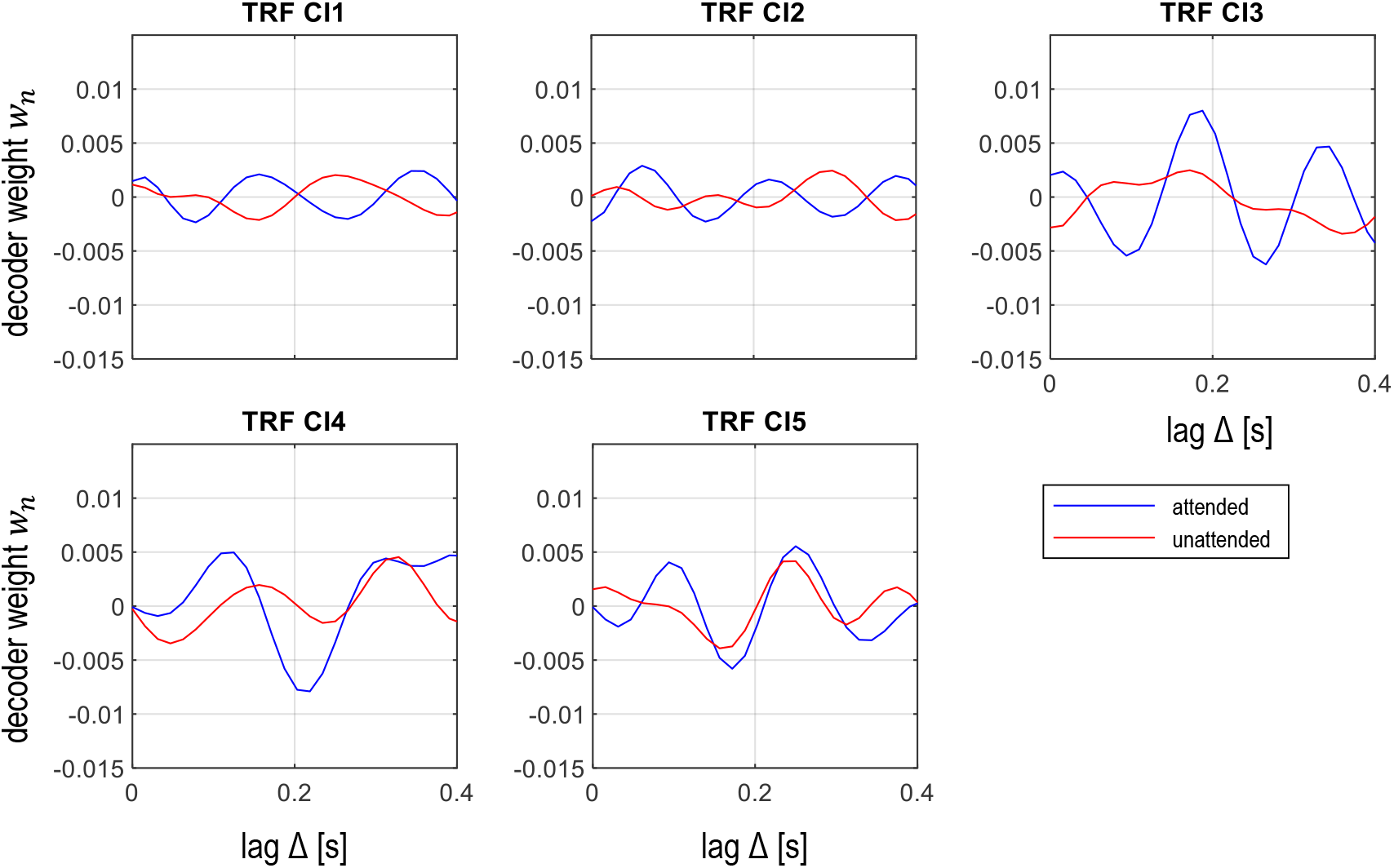
Temporal response functions (TRFs) of cochlear implant users (CI1-CI5) recorded through implant-based electroencephalography (iEEG). The linear forward model was trained with the envelope of the attended speech (blue) or the envelope of the unattended speech (red). The encoder weights *w*_*n*_ for the single-channel iEEG (case – E0) are shown across lag Δ between stimulus and recording.

### 3.4 Selective Attention Backward Decoding

#### 3.4.1 Optimization of Regularization Parameter λ and Lag Δ

The regularization parameter λ and the lag Δ both have a high impact on the decoding performance of the model. We evaluated the influence of λ and Δ on the decoding accuracy for both EEG modalities. Figures 8 and 9 show the decoding accuracies across Δ (from 0 to 480 ms) for different values of λ (0.001, 0.01, 0.1, 1, 10, and 100). Note that the chance level for EEG, estimated through a binomial test, ranged between 38.6 and 61.4 % (see Methods), while for iEEG it differed between the subjects and was thus not depicted in Figure 9.

**Figure 8:**
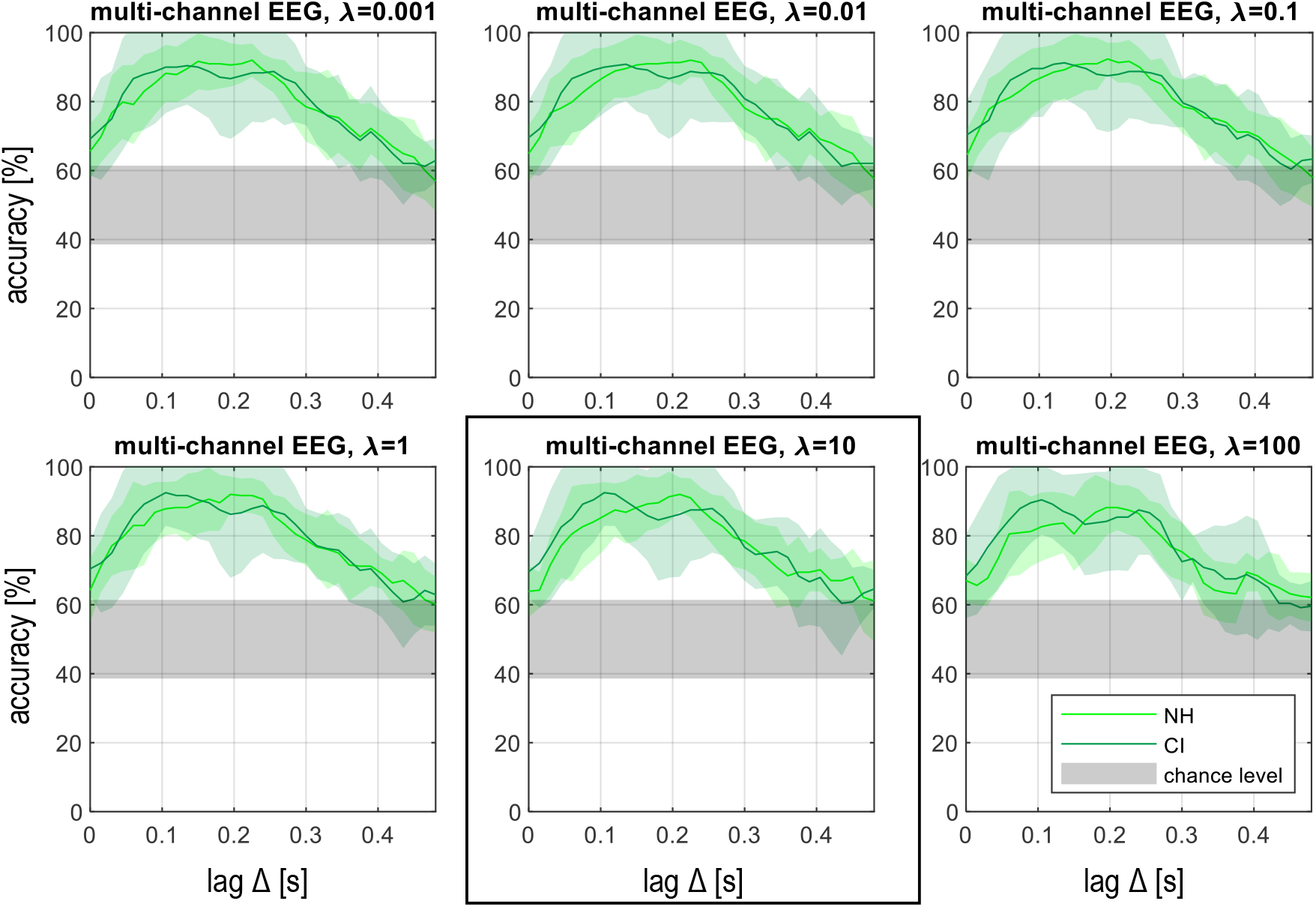
Mean decoding accuracies across 6 normal hearing (NH) listeners (light green) and 5 cochlear implant (CI) users (dark green) for all scalp electroencephalography (EEG) channels across lags Δ for different values of the regularization parameter *λ*. The colored shaded area denotes the standard deviation across subjects and the grey area denotes the chance level. The highest decoding accuracies (NH: 92.0 %; CI: 92.5 %) were achieved with *λ* = 10 with lags Δ of 210-225 ms and with *λ* = 10 with Δ of 105-120 ms.

**Figure 9:**
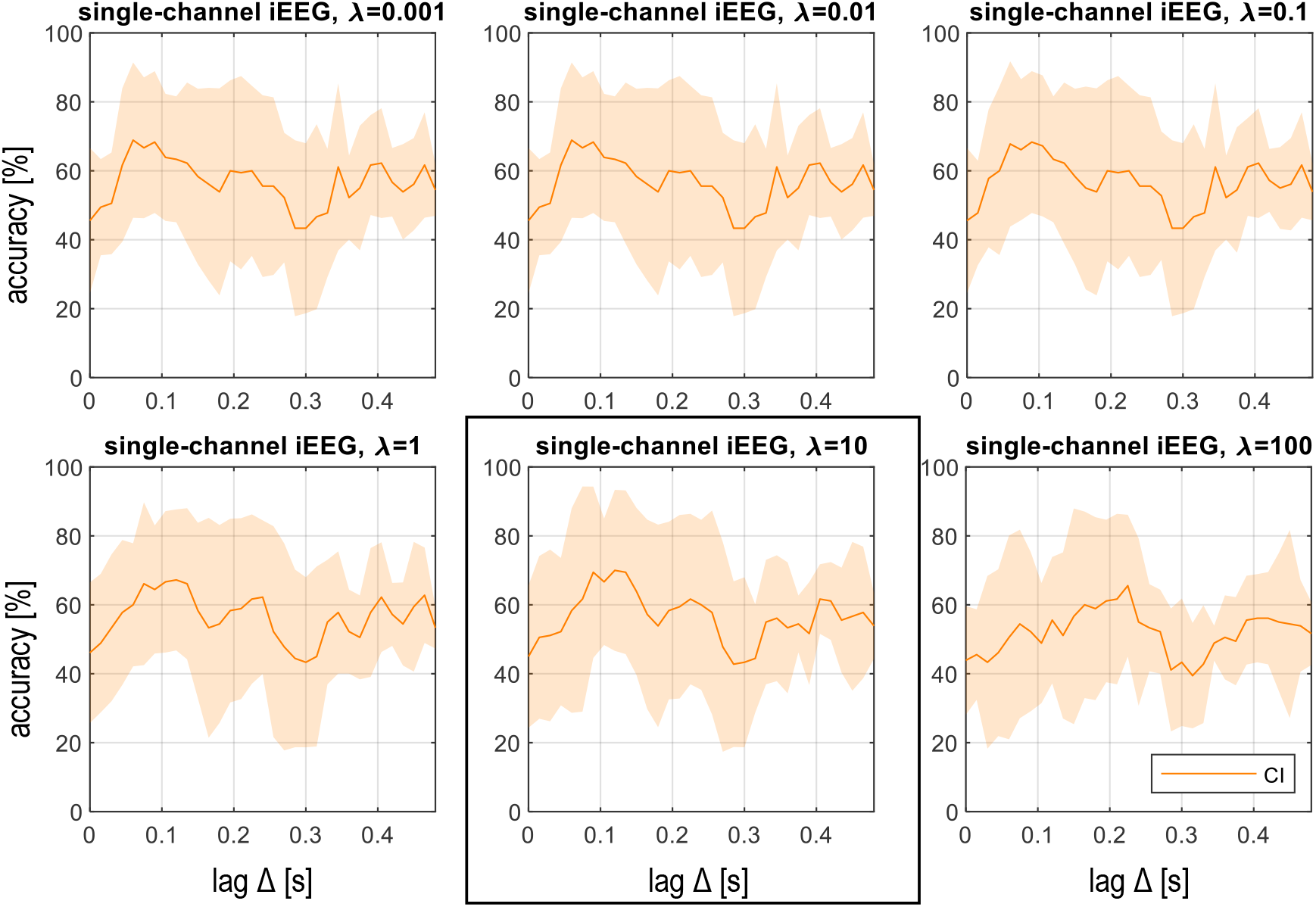
Mean decoding accuracies across 5 cochlear implant (CI) users for CI electrodes (case - E0) across lags Δ for different values of the regularization parameter *λ*. The shaded area denotes the standard deviation across subjects. The highest decoding accuracy of 70 % was achieved with *λ* = 10 and lags Δ of 120-135 ms. iEEG = implant-based electroencephalography.

For scalp EEG and iEEG, the highest decoding accuracies were obtained using a *λ* of 10. For scalp EEG the optimal Δ ranged between 105 and 210 ms. Note that for multi-channel and single-channel EEG we determined the same optimal *λ* for CI users, thus we present here only the data for multi-channel EEG. For iEEG the optimal Δ was 120 ms. These parameters (summarized in Table 3) were used in the following analyses for further comparisons between EEG and iEEG.

**Table 3:** Lag Δ that corresponds to the highest decoding accuracy across 5 cochlear implant (CI) users and 6 normal hearing (NH) listeners for different recordings. EEG = electroencephalography; iEEG = implant-based EEG.

|  | Multi-channel EEG | Single-channel EEG | Single-channel iEEG |
| --- | --- | --- | --- |
| CI | 105 ms | 105 ms | 120 ms |
| NH | 210 ms | 105 ms | - |

#### 3.4.2 Selective Attention Correlation Coefficient ρ Using Optimized Regularization Parameter λ and Lag Δ

We determined the Pearson correlation coefficient between the envelopes of the attended or the unattended speech and the reconstructed signal. The mean Pearson correlation coefficients ρ for all NH listeners and all CI users across lags Δ are shown in Figure 10.

**Figure 10:**
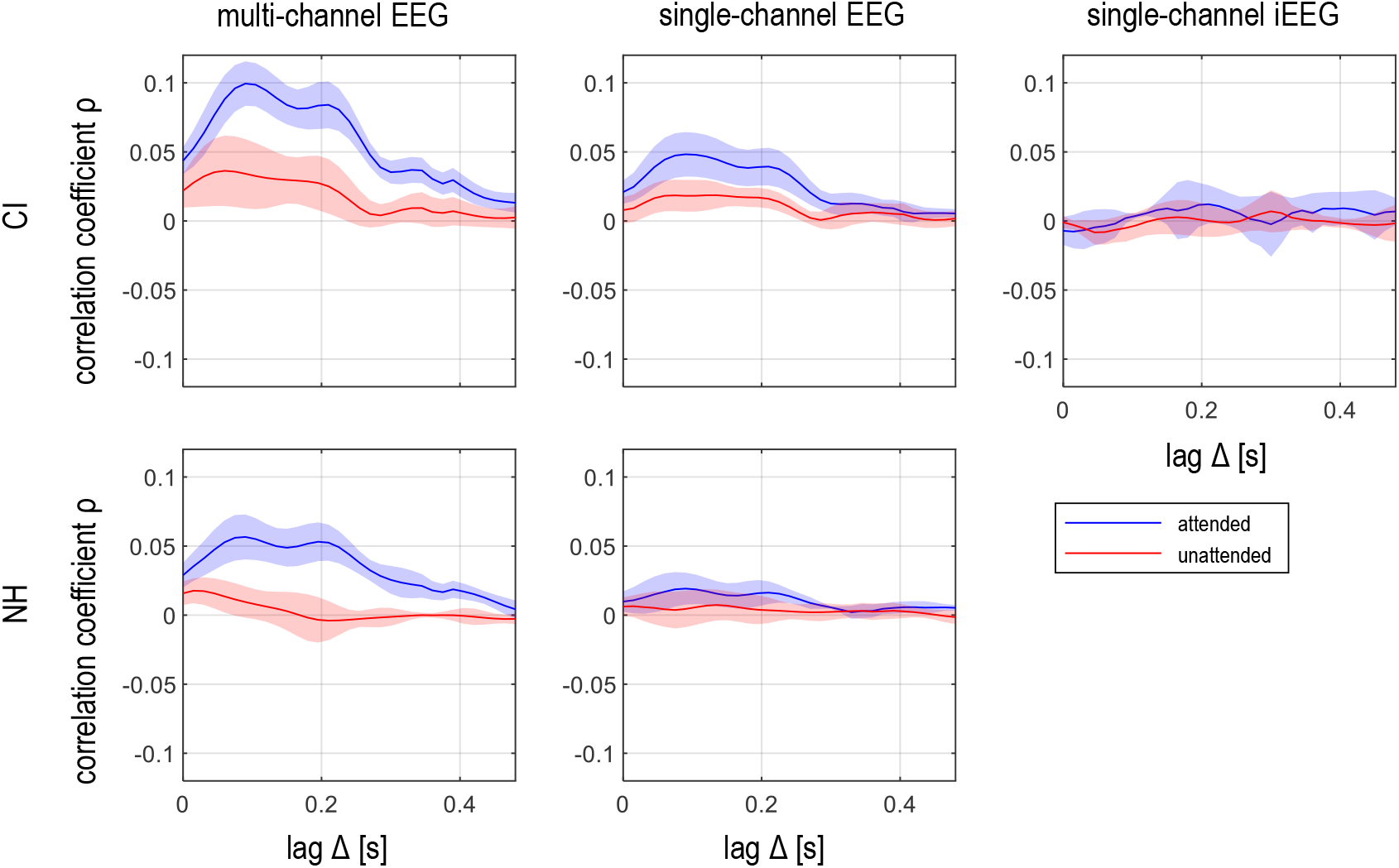
Mean Pearson correlation coefficients for 6 normal-hearing (NH) listeners (top) and 5 cochlear implant (CI) users (bottom) for multi-channel electroencephalography (EEG), single-channel EEG and single-channel implant-based EEG (iEEG). In each subplot the blue lines represent the correlation coefficient *ρ*_a_ between the reconstructed and the original envelope of the attended speech stream and the red lines represents the correlation coefficient *ρ*_u_ between the reconstructed and the original envelope of the unattended speech stream. The shaded area denotes the standard deviation across subjects.

The correlation coefficients for EEG present two distinct peaks at lags Δ = 100 ms and Δ = 200 ms. These lags correspond well to the peak latencies observed in the CAEPs (Figure 5) and the TRFs (Figure 6). The correlation coefficients for iEEG only presented a single peak at around Δ = 200 ms. Moreover, the attended correlation coefficient for iEEG was higher than the unattended correlation coefficient for the unattended speech at most lags.

The statistical analysis showed that the correlation coefficients ρ were not normally distributed, as assessed by the Shapiro-Wilk-Test (*p* < 0.05). The non-parametric Wilcoxon signed rank test showed that the correlation for the attended speech stream was significantly higher than for the unattended speech stream in all subjects with multi-channel EEG, in 8 out of 11 subjects with single-channel EEG and in 2 out of 5 subjects with iEEG. The details of the statistical analysis are provided in Table 4.

**Table 4:**
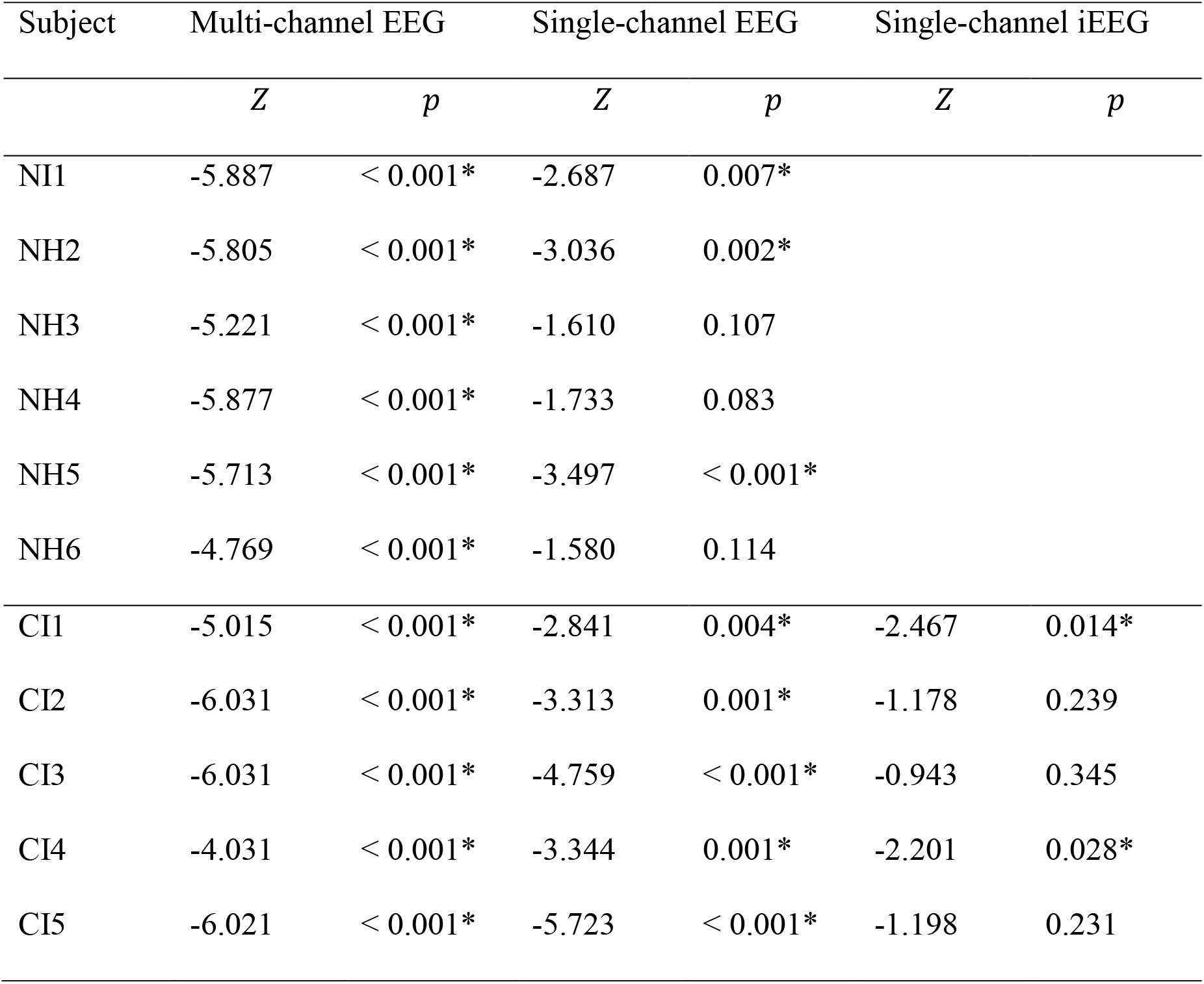
Results of the Wilcoxon signed rank test comparing the correlation coefficients *ρ* for the attended and for the unattended speech stream for optimized parameters. CI = cochlear implant; EEG = electroencephalography; iEEG = implant-based EEG; NH = normal hearing; ^*^ = significant results.

| Subject | Multi-channel EEG |  | Single-channel EEG |  | Single-channel iEEG |  |
| --- | --- | --- | --- | --- | --- | --- |
| | $Z$ | $p$ | $Z$ | $p$ | $Z$ | $p$ |
| NI1 | -5.887 | < 0.001* | -2.687 | 0.007* |  |  |
| NH2 | -5.805 | < 0.001* | -3.036 | 0.002* |  |  |
| NH3 | -5.221 | < 0.001* | -1.610 | 0.107 |  |  |
| NH4 | -5.877 | < 0.001* | -1.733 | 0.083 |  |  |
| NH5 | -5.713 | < 0.001* | -3.497 | < 0.001* |  |  |
| NH6 | -4.769 | < 0.001* | -1.580 | 0.114 |  |  |
| CI1 | -5.015 | < 0.001* | -2.841 | 0.004* | -2.467 | 0.014* |
| CI2 | -6.031 | < 0.001* | -3.313 | 0.001* | -1.178 | 0.239 |
| CI3 | -6.031 | < 0.001* | -4.759 | < 0.001* | -0.943 | 0.345 |
| CI4 | -4.031 | < 0.001* | -3.344 | 0.001* | -2.201 | 0.028* |
| CI5 | -6.021 | < 0.001* | -5.723 | < 0.001* | -1.198 | 0.231 |

#### 3.4.3 Selective Attention Decoding Accuracies Using Optimized Regularization Parameter *λ* and Lag Δ

The decoding accuracies with optimized parameters show a high inter-subject variability. A comparison of all subjects is shown in Figure 11 and the detailed statistics are provided in Table 5. With multi-channel scalp EEG, NH listeners and CI users reached 92.0 % and 92.5 % mean decoding accuracies, respectively. All subjects obtained accuracies above chance level. Using single-channel scalp EEG the performance dropped to 65.6 and 75.8 % for NH listeners and CI users, respectively. 9 out of 11 subjects obtained accuracies above chance level. Using the CI electrodes as sensors, the decoding accuracy dropped to 70.0 % in CI users but was still above chance level for 3 out of 5 subjects.

**Table 5:**
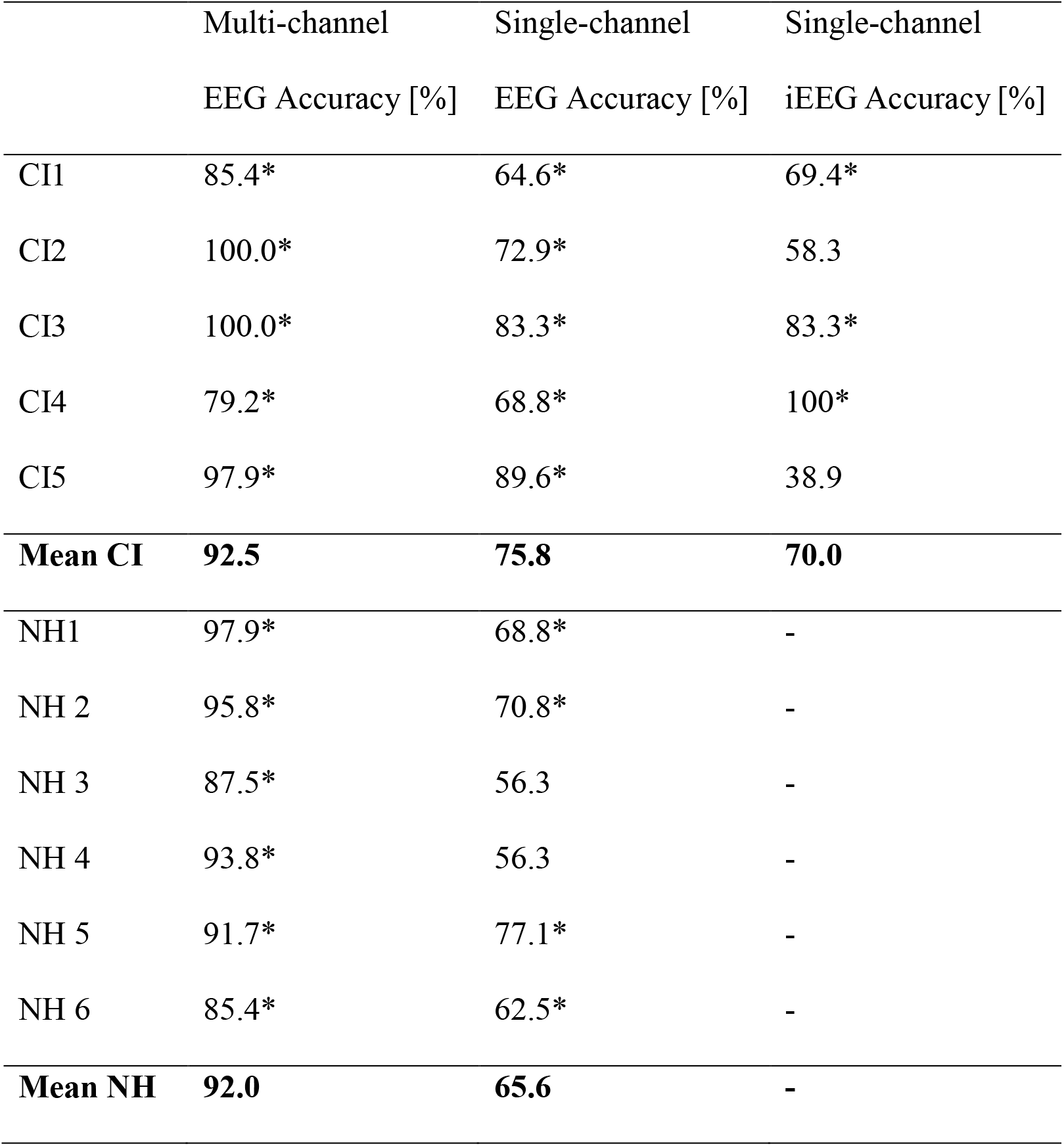
Individual and group mean decoding accuracy for normal hearing (NH) listeners and cochlear implant (CI) users using the best lag Δ and best regularization parameter *λ*. ^*^Accuracies that are above the subjects’ individual chance level. EEG = electroencephalography; iEEG = implant-based EEG.

**Figure 11:**
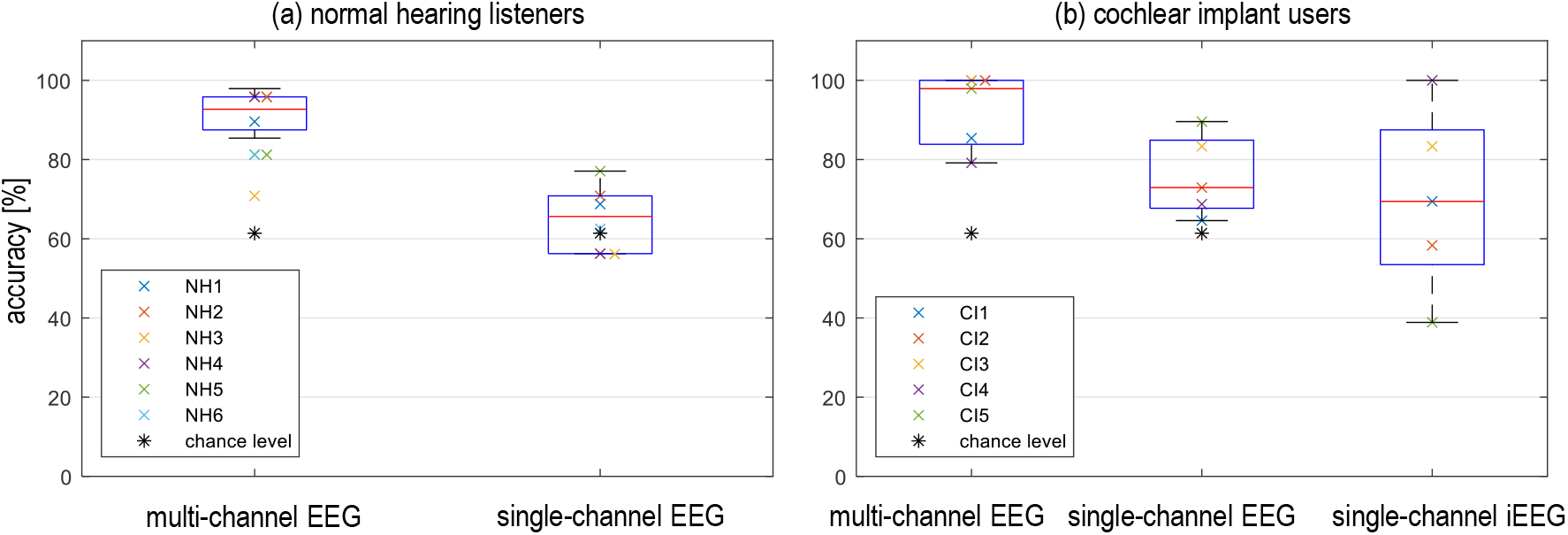
Individual decoding accuracies for the best lag Δ and best regularization parameter *λ* for (a) normal hearing listeners and (b) cochlear implant users; EEG = electroencephalography; iEEG = implant-based electroencephalography.

## 4. Discussion

The present study demonstrates the feasibility of recording CAEPs with the CI electrodes in combination with the CI backward telemetry system. Moreover, it demonstrates that selective attention can be decoded from EEG using such a CI-based recording. These findings might pave the way towards the development of closed-loop CIs.

### 4.1 Cortical Auditory Evoked Potentials (CAEPs)

Using the CI to record cortical potentials is a relatively unexplored field of research because the measurement requires long recording windows in the range of up to 1 s and most currently available systems are limited to short measurement windows due to hardware restrictions. McLaughlin et al. (2012) used the CI telemetry system from the company Cochlear Ltd. to measure CAEPs through repeated measurements concatenating EEG from time-shifted windows. In contrast to the present study, they used an electrode attached to the implant case of the CI (MP2) in combination with the most basal intracochlear electrode to record AEPs. They found N1 peaks in 3 CI users participating in the study but did not observe P2 peaks. Consistent to the results presented in the present study, they found that N1 was negative when the MP2 electrode on the implant case was referenced to an intracochlear electrode. The disadvantage of the approach of McLaughlin et al. (2012) was that they required the concatenation of the time-shifted windows causing extensive measurement time to record CAEPs. Somers et al. (2021) performed continuous EEG recording via a CI from the company Cochlear Ltd. using a percutaneous connector and a high-quality external amplifier. They tested different electrode configurations and found that the best configuration was a hybrid recording in which the Cz scalp electrode was referenced to an intracochlear electrode. They were also able to record CAEPs using only CI electrodes, even though the morphology was less clear and peaks were not detectable in all subjects. The CAEPs recorded in the present study show very similar morphologies to the CAEPs presented in the studies of McLaughlin et al. (2012) and Somers et al. (2021). However, the present experimental setup was not restricted by extensive measurement time and it was based on a clinical transcutaneous implant without requiring an additional external amplifier. A limitation of all studies is the small sample size. McLaughlin et al. (2012) and Somers et al. (2021) each included 3 subjects and the present study included 5 subjects. Further studies with more subjects are required in order to increase the robustness of the results to draw more general conclusions about the optimal electrode configuration.

### 4.2 Selective Attention Decoding

The CI-based recording technology offers the possibility to record near-to continuous cortical responses in an auditory selective attention paradigm. The present study showed that selective attention decoding from scalp EEG is feasible, even if the number of recording electrodes is highly reduced. Previous selective attention studies in NH listeners (e.g. Mirkovic et al., 2015; O’Sullivan et al., 2015) and CI users (Nogueira et al., 2019a, 2019b; Paul et al., 2020) focused on high-density scalp EEG with multiple recording channels. Mirkovic et al. (2015) analyzed the impact of reducing the number of electrodes and found that with a minimum of 25 electrodes the selective attention decoding performance in NH listeners remained stable. With five scalp EEG electrodes, they achieved a mean decoding accuracy of 74.1 % and with one channel the performance was 65 %. Using ear-EEG, which consists of 10 electrodes placed around each ear, in NH listeners Mirkovic et al. (2016) and Nogueira et al. (2019b) achieved a decoding accuracy of 70 % and 60 %, respectively. Fiedler et al. (2017) used one in-ear electrode in combination with one fronto-temporal scalp electrode to demonstrate that selective attention was possible using single-channel EEG. The mean selective attention decoding accuracy was 70 % and it was above chance level for the 7 NH listeners that participated in the study. The present study confirms the results from Fiedler et al. (2017) by showing that selective attention decoding was possible with single-channel scalp EEG. The mean selective attention decoding accuracy of 65.6 and 75.8 % for NH listeners and CI users, respectively, was in the same order of magnitude as the decoding accuracy observed in previous studies (Mirkovic et al., 2015; Fiedler et al., 2017). This result may be explained by the findings of Mirkovic et al. (2016) indicating that the number of recording channels might contribute less to the overall decoding performance than the electrode positions.

In the present study, a scalp vertex-to-mastoid electrode configuration was used. This configuration is optimal to capture the vertically orientated dipole arising from auditory cortices that mainly contributes to the early N1 and the P2 (e.g. Hine and Debener, 2007; Scherg and Von Cramon, 1986). Moreover, this vertex-to-mastoid configuration is supported by the location of the maximum weights of the topographic maps of selective attention TRFs (Crosse et al., 2016), which coincide with the vertex of the head, as also shown in Figure 6. Regarding the iEEG, the fixed implant case and intracochlear electrode positions limited the area and the orientation of the recording. However, the implant case electrode was located superior and lateral to the intracochlear electrodes. Thus, this configuration could capture both the vertically and radially oriented dipoles emanating from the auditory cortex (Scherg and Von Cramon, 1986). However, the exact electrode location could be affected by anatomical differences across subjects and differences in their CI implantation procedure. Further studies are needed to evaluate the effects of electrode position to suggest the most appropriate electrode configuration.

Previous selective attention studies with NH listeners (e.g. Mirkovic et al., 2015; O’Sullivan et al., 2015) and CI users (Nogueira et al., 2019a, 2019b; Paul et al., 2020) used a dichotic stimulus presentation in which one speech stream was presented to one ear and the second speech stream was presented to the contralateral ear. In contrast, the present study presented both speech streams monaurally to the same ear. This configuration may have degraded the selective attention performance of the subjects due to missing spatial separation between the talkers. However, the results of the present study indicate that selective attention decoding was possible in a monaural listening task. The mean decoding accuracy of 92.0 % for NH listeners was slightly better than similar studies who reported a decoding accuracy of up to 89 % in NH listeners using a linear backward model with a dichotic stimulus presentation (O’Sullivan et al., 2015).

The lower quality of the iEEG compared to the EEG may be explained by the internal electronics of the CI backward telemetry system, including the CI amplifier, the bit depth used by the telemetry system as well as other error sources. The CI amplifier runs on low power and therefore provides lower gain than a conventional EEG amplifier. In addition, the drift in the iEEG data caused signal saturation after a variable period of time between 15 and 90 s. The time at which saturation occurred depends on individual characteristics of the CI user, including the skin thickness which may influence the radiofrequency connection of backward telemetry system. The minimum possible recording time of 15 s was already longer than the recording times used in previous studies to record CAEPs (McLaughlin et al., 2012; Somers et al., 2021). However, to obtain more reliable data for selective attention decoding in all subjects, the recording time needs to be further extended or the paradigm needs to be adjusted accordingly in a future study.

### 4.3 Future Outlook and Potential Applications

The use of CI electrodes in combination with the CI backward telemetry system to record cortical potentials is a promising method with multiple future possibilities. In order to be applicable in the future, stimulation and recording will need to be performed simultaneously. Because of the close proximity between stimulating and recording electrodes, a high contamination by stimulation artifact can be expected. Since common EEG artifact rejection techniques, such as independent component analysis (ICA), are not suitable for single-channel recording, new strategies will need to be explored. An example is presented in the study by Somers et al. (2018) who included small gaps within the stimulation and recorded EEG only in those artifact-free gaps.

The advantage of a CI-based EEG in comparison to a conventional scalp EEG is that no additional hardware is needed and that it is easily applicable in a clinical environment or in daily life. The technology could be used to develop a closed-loop CI with neuro-feedback functionality. Such a closed-loop CI could be used for objective fittings in children or disabled people who cannot provide adequate feedback. It could also be used for long-term monitoring of CI users by continuous recording of AEPs or selective attention decoding. Ultimately, the goal of closed-loop CIs would be to provide assistance in challenging listening environments, such as cocktail party scenarios. A combination of selective attention decoding algorithms, as presented in the present study, and source separation algorithms for CIs (e.g. Goehring et al., 2017; Tahmasebi et al., 2020) could improve speech performance of CI users. Recently, Ceolini et al. (2020) showed that brain-informed speech separation (BISS) improves speech separation in the front end of hearing aids. Thus, neuro-steered CIs or hearing aids that adapt sound processing through attention show a high future potential.

## 5. Conclusions

The present study confirms that it is possible to decode selective attention in NH and hearing impaired listeners in a monaural listening task. The selective attention decoding was possible using multi-channel and also using single-channel high-density scalp EEG, even though the decoding accuracy with single-channel data was lower. Using CI electrodes and a CI backward telemetry system to record electrophysiological signals remains a challenge. In this work, the general feasibility of using CI-based recordings to capture CAEPs and to decode selective attention has been shown, however selective attention decoding with CI-based recordings was lower than with scalp EEG. Despite a high inter-subject variability it was possible to record CAEPs in 2 out of 5 CI users with contralateral acoustic hearing and to decode selective attention in 3 out of 5 CI users with contralateral acoustic hearing through CI-based recordings.

## Acknowledgments

We would like to thank Hanna Dolhopiatenko and Anna Ruhe for their help during the conduction of the experiments. Moreover we thank all participants for their time and patience. We would like to thank Kanthaiah Koka, Chen Chen and colleagues from Advanced Bionics for providing research tools and advice. This work was supported by the Deutsche Forschungsgemeinschaft (DFG, German Research Foundation) under Germany’s Excellence Strategy - EXC 2177 - Project ID 390895286.

## Ethical statement

The study was carried out in accordance with the declaration of Helsinki principles and approved by the ethics committee of the Hannover Medical School (Hanover, Germany). The participants provided their written informed consent to participate in this study.

*This is the version of the article before peer review or editing, as submitted by the authors to the Journal of Neural Engineering. IOP Publishing Ltd is not responsible for any errors or omissions in this version of the manuscript or any version derived from it*.

